# Combined Molnupiravir and Nirmatrelvir Treatment Improves the Inhibitory Effect on SARS-CoV-2 in Rhesus Macaques

**DOI:** 10.1101/2022.09.03.506479

**Authors:** Kyle Rosenke, Matt C. Lewis, Friederike Feldmann, Eric Bohrnsen, Benjamin Schwarz, Atsushi Okumura, W. Forrest Bohler, Julie Callison, Carl Shaia, Catharine M. Bosio, Jamie Lovaglio, Greg Saturday, Michael A. Jarvis, Heinz Feldmann

## Abstract

The periodic emergence of SARS-CoV-2 variants of concern (VOCs) with unpredictable clinical severity and ability to escape preexisting immunity emphasizes the continued need for antiviral interventions. Two small molecule inhibitors, molnupiravir (MK-4482), a nucleoside analog, and nirmatrelvir (PF-07321332), a 3C-like protease inhibitor, have each recently been approved as monotherapy for use in high risk COVID-19 patients. As preclinical data are only available for rodent and ferret models, we originally assessed the efficacy of MK-4482 and PF-07321332 alone and then in combination Against infection with the SARS-CoV-2 Delta VOC in the rhesus macaque COVID-19 model. Notably, use of MK-4482 and PF-07321332 in combination improved the individual inhibitory effect of both drugs. Combined treatment resulted in milder disease progression, stronger reduction of virus shedding from mucosal tissues of the upper respiratory tract, stronger reduction of viral replication in the lower respiratory tract, and reduced lung pathology. Our data strongly indicate superiority of combined MK-4482 and PF-07321332 treatment of SARS-CoV-2 infections as demonstrated here in the closest COVID-19 surrogate model.

**One Sentence Summary:** The combination of molnupiravir and nirmatrelvir inhibits SARS-CoV-2 replication and shedding more effectively than individual treatments in the rhesus macaque model.

## Introduction

Severe acute respiratory syndrome coronavirus 2 (SARS-CoV-2), the causative agent of coronavirus disease 2019 (COVID-19) [1], was initially reported in China in December 2019 [2] and has subsequently spread through the rest of Asia and around the World. As of writing (August 2022), the pandemic accounts for over 599 million confirmed SARS-CoV-2 infections and 6.47 million deaths. Following an initial relatively stable phase, the pandemic is now being driven by regional and global waves of newly emerging SARS-CoV-2 variants. These variants emerge on a steady but unpredictable basis. To date, 13 variants of interest (VOIs) or concern (VOCs) associated with mutations that alter key aspects of transmissibility, immune evasion or severity of disease have emerged, excluding the 7 Omicron subvariants currently circulating [3, 4]. Mutations associated with VOCs are most frequently located within the spike protein where they affect functions such as receptor binding affinity and antibody neutralization capacity but can be located throughout the entire SARS-CoV-2 genome [3–9].

Molnupiravir (MK-4482) and nirmatrelvir (PF-07321332) are orally administered antivirals that have recently been approved for use to treat high risk COVID-19 patients [10–13]. These compounds target distinct stages of the SARS-CoV-2 replication cycle, but neither directly targets stages involving the viral spike protein [14, 15]. MK-4482 is a nucleoside analogue affecting the SARS-CoV-2 polymerase fidelity resulting in a catastrophic rate of mutation that ultimately reduces progeny virus infectivity [15]. PF-07321332 is a 3C-like protease inhibitor that prevents cleavage of the SARS-CoV-2 polyprotein, again ultimately inhibiting viral replication [14]. PF-07321332is currently used in combination with low dose ritonavir, an inhibitor of cytochrome P450-3A4, where ritonavir increases the PF-07321332 serum half-life allowing for increased potency [14]. MK-4482 is used under the trade name Lagevrio™. PF-07321332 together with ritonavir comprises the commercial COVID-19 treatment, Paxlovid™.

Results from recent studies in tissue culture and rodent and ferret models suggested that both MK-4482 and PF-07321332 retain activity when used as monotherapies against emerging VOCs including Omicron [16–18]. Herein, we originally investigated the efficacy of MK-4482 and PF-07321332, both as individual monotherapies and in combination, against the SARS-CoV-2 Delta VOC in the rhesus macaque COVID-19 model [19]. Notably, use of MK-4482 and PF-07321332 in combination improved the individual inhibitory effect of both drugs on clinical outcome, viral RNA and infectious viral loads in the upper and lower respiratory tract of infected animals. Our study demonstrates efficacy of MK-4482 and PF-07321332 as monotherapy, but with improved efficacy when used in combination, against SARS-CoV-2 in the closest COVID-19 surrogate model.

## Results

### Study design

To assess the efficacy of the two treatments individually and in combination, rhesus macaques were randomly divided into vehicle- or MK-4482-, PF-07321332- or combination treatment groups (n=5 per group). Animals were infected with 2×10^6^ TCID_50_ of SARS-CoV-2 Delta VOC by combined intranasal and intratracheal routes (1×10^6^ TCID_50_ each route). Drug treatments began 12 hours post-infection (12hpi) with animals receiving either vehicle or 130mg/kg MK-4482 (260mg/kg/day) or 20mg/kg PF-073211332 + 6.5mg/kg ritonavir (40mg/kg/day, 13mg/kg/day) or a combination of all 3 compounds consisting of the same doses as for individual treatments. Treatments were administered by oral gavage every 12h (7 treatments in total). The study ended 4 days post-infection (dpi) (12h following last treatment), at which time animals were euthanized for tissue collection and analysis (Fig. 1A).

**Figure 1.**
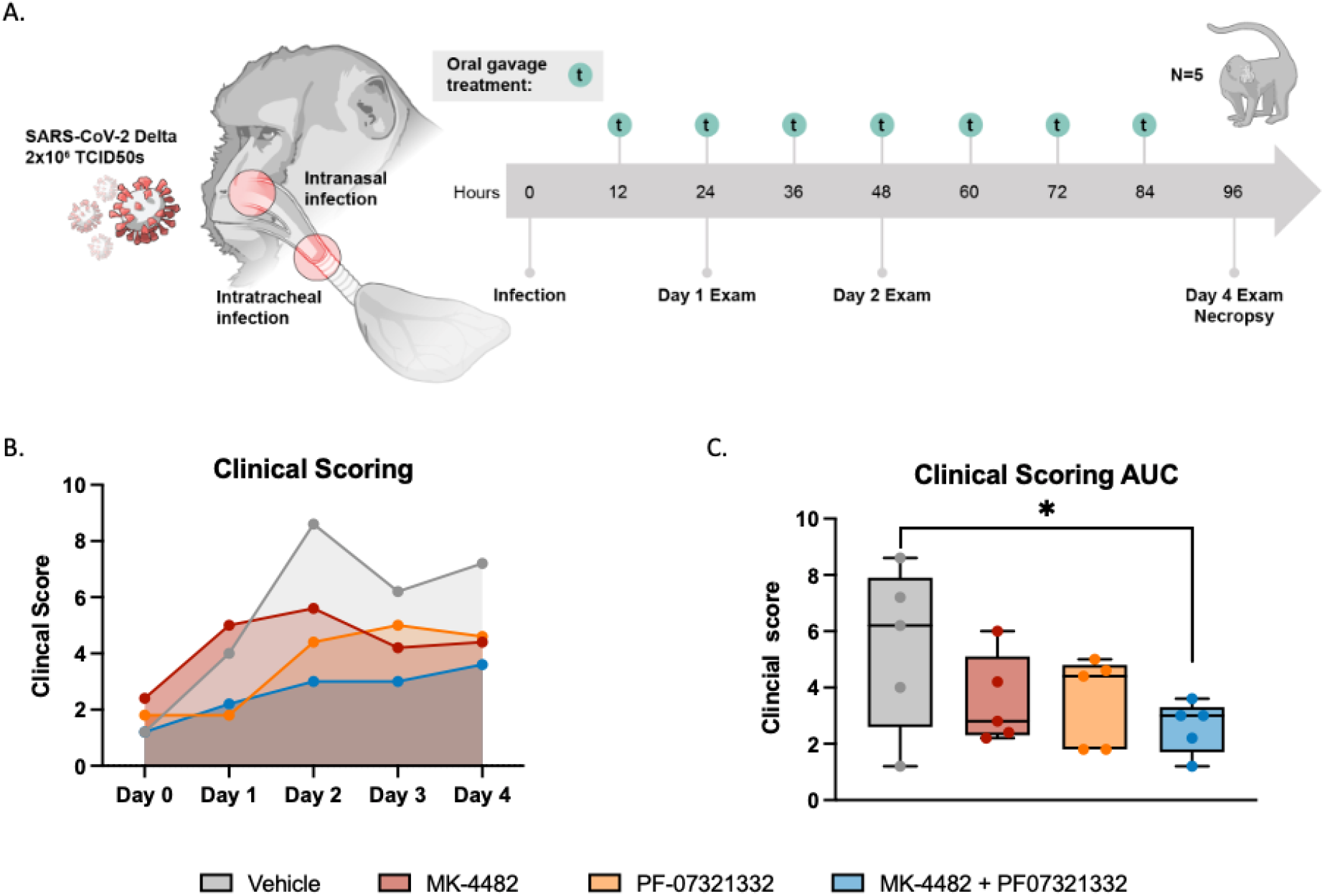
Combination therapy of MK-4482 and PF-07321332 reduced clinical scoring in rhesus macaques. **Experimental design (A)**. Rhesus macaques (n = 5) were infected with 2 × 10^6^ TCID_50_ SARS-CoV-2 by combined intranasal and intratracheal routes. Treatments were started 12 hours post-infection, with continued dosing every 12 hours. Clinical exams were conducted on 1, 2 and 4 dpi and animals were euthanized and necropsied on 4dpi. **Clinical scoring (B)**. Animals were scored daily for clinical signs of disease over the course of the study. **AUC analysis of clinical scoring (C)**. Clinical scores were calculated for each animal per day, AUC was then calculated over the course of the study and displayed in minimum-to-maximum boxplot with median displayed. Ordinary one-way ANOVA with multiple comparisons were used to evaluate significance (*P-value = 0.01 to 0.05).

### Combination treatment results in a significant reduction in clinical disease

Animals were scored for signs of disease daily by the same person blinded to the study groups using a previously established scoring sheet [19]. A score (0–15) was assigned for each of the following: general appearance, skin and fur, nose/mouth/eyes/head, respiration, feces and urine, food intake, and locomotor activity. All scoring was performed prior to anesthesia for treatments and examinations. Groups scored evenly in the morning prior to inoculation (0dpi). Animals in the vehicle treated group showed the highest scores throughout the experiment peaking at 2dpi. Animals in the group receiving the combination treatment scored the lowest throughout (Fig. 1B). Animals in the groups treated with either MK-4482 or PF-07321332 alone scored in between (Fig. 1B). Although differences between groups were not statistically significant on any single day of the study, AUC analysis showed a significant difference between the vehicle treated group and the group receiving the combination therapy (Fig 1C).

### Combination treatment results in a significantly greater reduction of SARS-CoV-2 shedding than monotherapy alone

Nasal and oral swabs were collected from rhesus macaques during examinations performed on 1, 2 and 4dpi. Subgenomic E (sgE)-based RT-PCR was used as an initial measure of active SARS-CoV-2 replication [20]. Compared to the vehicle treated group, sgE viral RNA loads in the nasal swabs were lower in all three treatment groups and remained significantly lower in the combination therapy group for the duration of the study. The PF-07321332, but not MK-4482, monotherapy treatment group was also significantly lower at 4dpi (Fig 2A). These differences were further amplified in the infectious titers recovered from the nasal swabs. At 1dpi, all treated animals had significantly lower infectious titers. By 2dpi, the differences between treated and untreated animals were further amplified and a larger subset of animals in each treatment group had no detectable infectious virus (Fig 2B). There were no significant differences in the levels of viral RNA detected in the oral swabs at any timepoint examined (Fig 2C), but AUC analysis performed for the entirety of the study did reveal a significant difference between the combination therapy and the vehicle control (Fig S1a). Infectious virus was significantly lower at 1dpi in all treated groups when compared to the vehicle control group, and treated groups remained significantly lower over the entirety of the study (Fig 1D).

**Figure 2.**
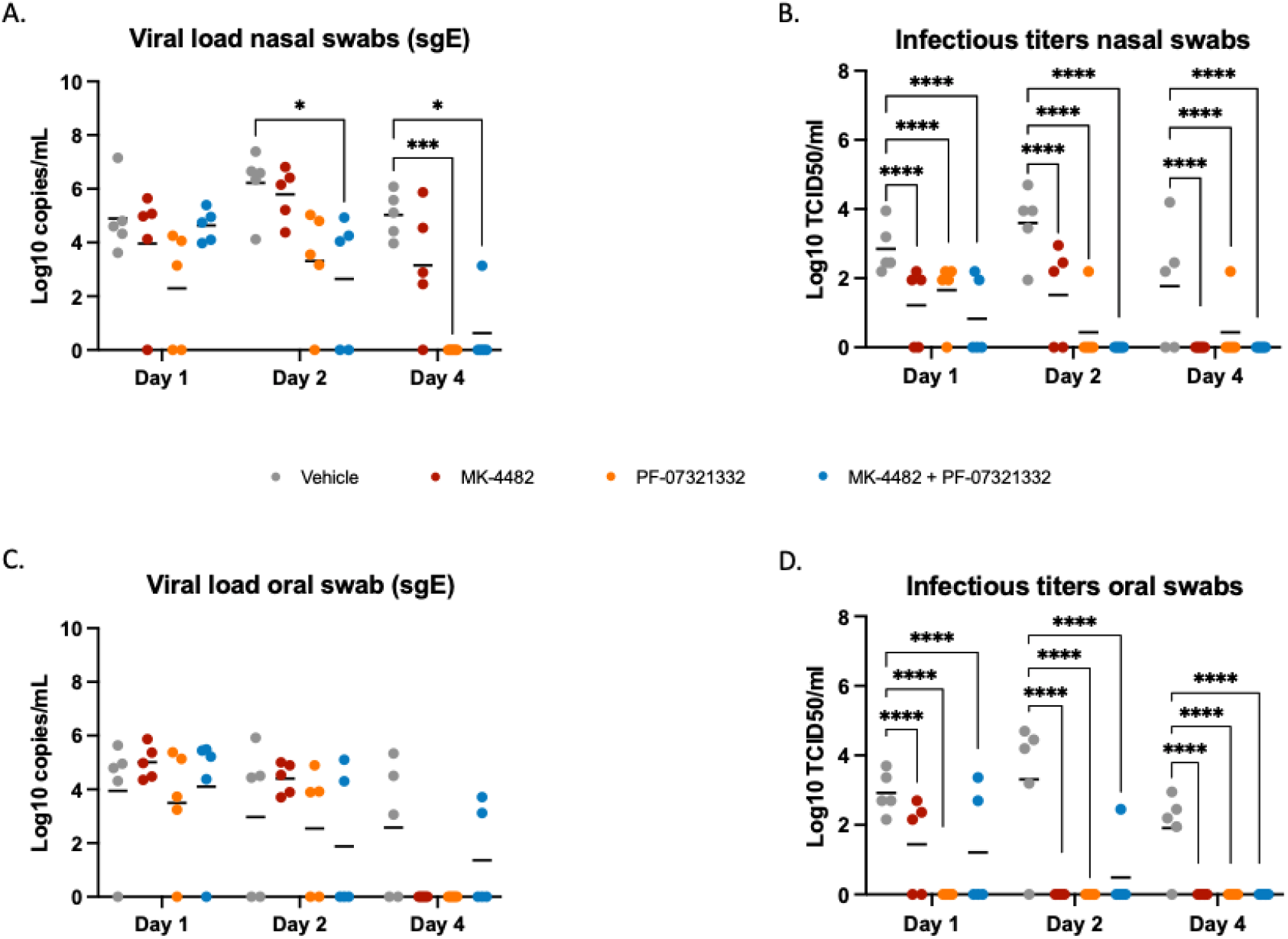
Combination therapy of MK-4482 and PF-07321332 significantly reduced virus replication in the upper respiratory tract of SARS-CoV-2 infected rhesus macaques. **Viral RNA load in nasal (A) and oral swabs (C)**. Nasal and oral swabs were collected on 1, 2 and 4dpi and viral RNA loads were determined by quantitative RT-PCR targeting sgE RNA. **Infectious titers in nasal (B) and oral (D) swabs**. Nasal and oral swabs were collected on 1, 2 and 4dpi and infectivity was determined by using a tissue culture infectious dose (TCID) assay with virus titers presented as TCID_50_/ml. Statistical differences in viral load and infectious virus titers in each study arm were assessed by a two-way ANOVA using Tukey’s multiple comparisons test (P-values, * = 0.01 to 0.05, ** = 0.001 to 0.01, *** = 0.0001 to 0.001, **** = <0.0001).

### Combination treatment results in a significantly greater reduction of SARS-CoV-2 replication in the lower respiratory tract

Bronchoalveolar lavages (BALs) were collected during each clinical examination at 1, 2 and 4dpi. As shown in Fig 3A, sgE RNA loads were lower in PF-07321332 and combination therapy groups at 1dpi and 2dpi, compared to the vehicle controls. The differences were more pronounced between groups at 2dpi but at no time was the sgE significantly different between groups (Fig 3A). AUC analysis of the BALs did reveal a significant difference between the vehicle treated and combination treated groups (Fig S1b). Infectious virus in the BAL samples were lower in all treatment groups at 1dpi and 2dpi as compared to the vehicle control group. The combination therapy was significantly lower than the vehicle controls at 1dpi (Fig. 3B). Although only one animal in the MK-4482 group and no animals in the combination therapy groups had detectable infectious virus at 2dpi, these results were not statistically significant. By 4dpi all groups had a single animal with detectable levels of infectious virus with all other animals being negative (Fig 3B).

**Figure 3.**
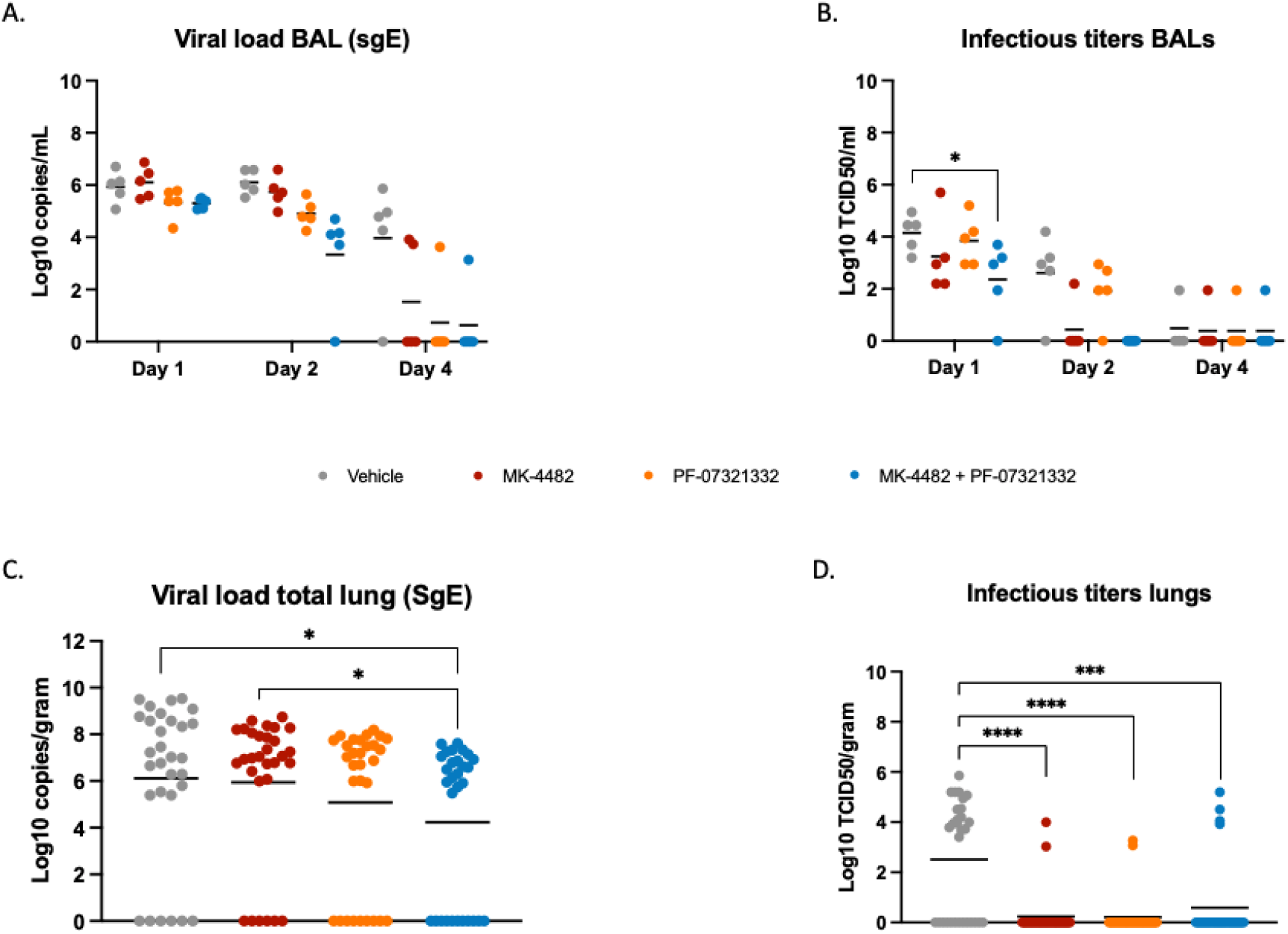
Combination therapy of MK-4482 and PF-07321332 significantly reduced viral replication in the lower respiratory tract of SARS-CoV-2 infected rhesus macaques. **Viral RNA load in BAL (A) and total lung lobes (C)**. BAL samples were collected on 1, 2 and 4dpi and lung samples were collected following necropsy on 4dpi. Viral RNA loads from BAL and lung samples were determined by quantitative RT-PCR targeting sgE RNA. **Infectious titers in BAL (B) and total lung lobes (D)**. BAL samples were collected on 1, 2 and 4dpi and lung samples were collected following necropsy on 4dpi. Infectivity was determined using a TCID_50_ assay and are presented as TCID_50_/ml. Samples from each lung lobe (n=6) of each animal were assessed individually and compiled for analysis **(C, D)**. Statistical differences in viral load and infectious virus titers in the BAL were assessed by two-way ANOVA using Tukey’s multiple comparisons test. Statistical differences in total lung lobes were analyzed with Kruskal-Wallis with Dunn’s multiple comparisons test (P-values, * = 0.01 to 0.05, ** = 0.001 to 0.01, *** = 0.0001 to 0.001, **** = <0.0001).

Tissue samples from each lung lobe were collected at 4dpi for viral load analysis. RNA was isolated from each sample for PCR analysis. Each lobe value was then pooled for total lung comparisons between groups. Animals receiving PF-07321332 and the combination therapy had lower levels of sgE RNA viral loads in the lungs than the vehicle treated group; the combination therapy group had significantly less detectable sgE viral loads than either the vehicle and MK-4482 treated groups (Fig 3C). Infectious virus loads in lungs were significantly reduced in all treatment groups by study end at 4dpi (Fig 3D).

### Treated animals had less pathology and viral antigen in the upper and lower respiratory tissues

Histological analysis of samples removed from each lung lobe revealed treatments decreased lung pathology. The vehicle controls developed minimal-to-marked pathology in four of five animals. These lesions were characterized as multifocal, mild to marked interstitial pneumonia characterized by thickening of alveolar septa by edema fluid and fibrin and small to moderate numbers of macrophages and fewer neutrophils. Alveoli contained increased numbers of pulmonary macrophages and neutrophils. Multifocal perivascular infiltrates of small to moderate numbers of lymphocytes forming perivascular cuffs were also observed. There was minimal type II pneumocyte hyperplasia consistent with the early SARS-CoV-2 disease progression (4dpi) (Fig. 4A). All treatment groups exhibited a decreased overall presence of interstitial pneumonia compared to the vehicle controls. MK-4482 treatments resulted in a reduction in observable interstitial pneumonia to 3 out of 5 animals and those lesions were minimal (Fig. 4B). Treatment with PF-07321332 reduced the number of effected animals to 2 of 5, with both scoring minimal and mild in one lung lobe each (Fig. 4C). Treatment with the drug combination reduced pneumonia even further with only 1 of 5 animals displaying minimal lesions in two lung lobes (Fig. 4D). Immunohistochemistry (IHC) of the same lung samples revealed a reduction in observable viral antigen between the vehicle treated group and all 3 treated groups (Fig 4E-H). Nasal turbinates were also collected at the time of euthanasia with no observable pathology in either the respiratory or olfactory epithelium (Fig S2). Although there was an absence of pathology, IHC analysis did show scattered immunoreactivity in the vehicle treated animals and little to no observable viral antigen in treated groups (Fig S2).

**Figure 4:**
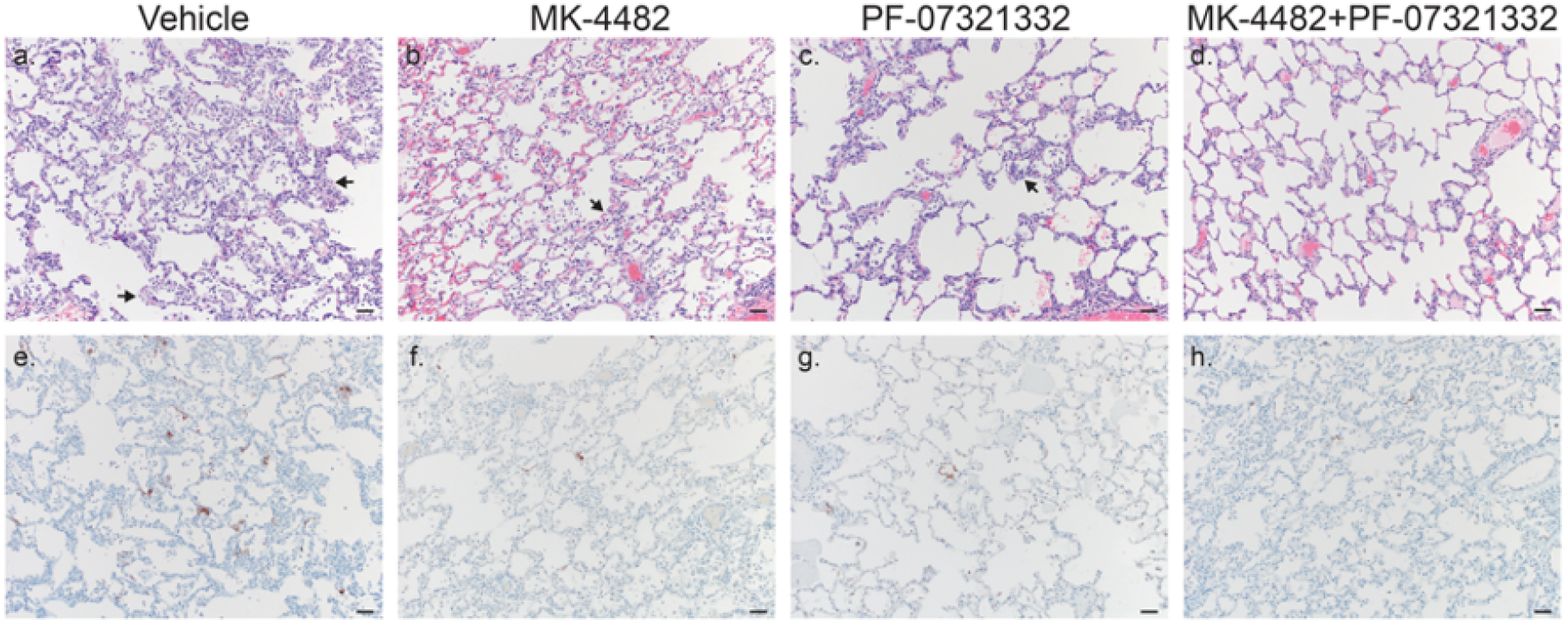
Combination therapy reduced lung pathology and SARS-CoV-2 antigen load in SARS-CoV-2 infected rhesus macaques. Lung tissues were collected on 4dpi and stained with either hematoxylin and eosin (H&E) or immunohistochemistry (IHC). **H&E staining of representative tissues sections of the lungs (A-D). IHC staining of SARS-CoV-2 antigen in corresponding representative lung sections (E-H). (A)** Vehicle control animals showed moderate interstitial pneumonia (arrows) and moderate immunoreactivity of type I pneumocytes in 4 of 5 animals **(A & E)**. MK-4482 treated animals showed mild interstitial pneumonia (arrows) and scattered immunoreactive type I pneumocytes in 3 of 5 animals **(B & F)**. PF-07321332 treated animals showed mild interstitial pneumonia (arrows) and scattered immunoreactive type I pneumocytes in 2 of 5 animals **(C & G)**. Combination treatment animals basically showed no interstitial pneumonia with 1 of 5 animals displaying minimal lesions and scattered immunoreactive type I pneumocytes **(D & H)**. 100X, Bar=100µm

### Bioavailable drugs were detected in both sera and lung samples

To assess circulating drug levels across each treatment regimen to ensure bioavailability in the pulmonary compartment, levels of MK-4482, PF-07321332, and ritonavir were quantified in plasma collected prior to drug administration at each clinical exam time point and in clarified lung homogenates collected at the time of necropsy (Table 1). As previously described [21], EIDD-1931, the active nucleobase metabolite of the MK-4482 pro-drug, was used as a surrogate for MK-4482; MK-4482 signals were analyzed in all samples and a standard curve was assessed but no signals above the limit of detection were observed as expected due to its rapid metabolism. As plasma was collected prior to dosing at each exam point, levels of each therapeutic molecule in the plasma reflect the lowest circulating concentrations over the treatment course. At each time point, levels of all 3 drugs were readily detectable, with a mean of 46.47nm for EIDD-1931 and 12.75nm for PF-07321332 in plasma. Drug concentrations were lower in lung tissue with a mean of 21.52nmol/g EIDD-1931 and 0.06nmol/g PF-07321332. Combination therapy increased these values in both plasma and lungs to 76.17nM EIDD-1931 and 19.21nM PF-07321332 in plasma and 23.84 nmol/g EIDD-1931 and 0.36 nmol/g PF-07321332 in the lungs. All values agree with the treatment scheme in both plasma and lung homogenate samples, with anticipated slight temporal fluctuations. Lung levels of EIDD-1931 were also in good agreement with previous data examining efficacy of MK-4482 in the Syrian golden hamster model [21].

**Table 1:** Measurements of pharmaceutical active or surrogate pharmaceutical metabolite in each treatment group. Measurements were made in plasma at each clinical examination point prior to dosing and in lung homogenate at the study end point from a single lobe of the lung. The EIDD-1931 metabolite was used as a surrogate for MK-4482 and its associated metabolites due to instability of MK-4482. The mean and standard error on the mean are displayed (plasma: n=15; lung tissue: n=5).

| Treatment | Vehicle | MK-4482 | PF-07321332 | MK-4482 + PF-07321332 |
| --- | --- | --- | --- | --- |
| **Plasma** |  |  |  |  |
| EIDD-1931 (nmol) | 0 | 46.47 ± 7.31 | 0 | 76.17 ± 15.53 |
| PF-07321332 (nmol) | 0 | 0 | 12.75 ± 4.41 | 19.21 ± 7.46 |
| Ritonavir (nmol) | 0 | 0 | 0.70 ± 0.36 | 0.31 ± 0.13 |
| **Lung** |  |  |  |  |
| EIDD-1931 (nmol/g) | 0 | 21.52 ± 7.00 | 0 | 23.84 ± 6.58 |
| PF-07321332 (nmol/g) | 0 | 0 | 0.06 ± 0.03 | 0.36 ± 0.08 |
| Ritonavir (nmol/g) | 0 | 0 | 0.03 ± 0.02 | 0.13 ± 0.06 |

## Discussion

SARS-CoV-2 VOCs, which are now driving the pandemic, are characterized by their altered transmission rate, pathogenicity or evasion from prior immunity in individuals either vaccinated or previously infected [3, 6]. This continued emergence and selection of SARS-CoV-2 variants within populations with preexisting levels of spike-focused immunity remains a threat to the long-term effectiveness of all current commercial SARS-CoV-2 vaccines, which are all spike-based, as well as spike-directed treatment approaches such as antibody therapy. It is therefore important to improve existing drugs/dosing regimens and to continue the development of new drugs that work independently from an effect on the spike protein. Antiviral compounds targeting SARS-CoV-2 replication and processing of the viral polyprotein are obvious candidates. MK-4482 and PF-07321332 are compounds targeting the SARS-CoV-2 polymerase and protease, respectively [14, 15]. Both compounds have shown SARS-CoV-2 antiviral efficacy as single treatments in tissue culture [18, 21] and rodent and ferret disease models [21–25]. Molnupiravir (MK-4482) and Paxlovid™ (PF-07321332 and ritonavir) have recently been approved for use in patients considered high risk for developing severe COVID-19 [11, 12].

Neither MK-4482 or PF-07321332 have been evaluated for their antiviral efficacy in a SARS-CoV-2 nonhuman primate (NHP) disease model. Using the rhesus macaque model of SARS-CoV-2 infection [19] we assessed the anti-viral activities of both compounds individually as well as in combination against the SARS-CoV-2 Delta VOC. The Delta VOC was selected as this variant has been shown to cause the most severe infection of all VOCs, especially compared to the rather mild infection by Omicron, tested in the rhesus macaque model to date [26]. Utilization of the more pathogenic Delta VOC therefore assured use of the most stringent NHP challenge model available to measure treatment efficacy. As shown here, individual treatment with both compounds resulted in significantly reduced SARS-CoV-2 viral load in the upper and lower respiratory tract significantly reducing SARS-CoV-2 shedding and replication, and ultimately led to less severe respiratory disease compared to vehicle treated animals.

As the mode of action for MK-4482 and PF-07321332 are distinct, the former being a nucleoside analog and the latter an inhibitor of viral protease inhibitor, combined therapy may provide a potential benefit over monotherapy. This was supported by recent *in vitro* data demonstrating a synergistic antiviral effect of MK-4482 and PF-07321332 against Delta and Omicron VOCs in comparison to individual drug treatment [27, 28], and *in vivo* for a Korean strain isolated early in the pandemic [29] using a transgenic mouse model [30]. In our present study, combined administration of MK-4482 and PF-07321332 treatment in the rhesus macaque model was well tolerated as indicated by clinical observation and blood chemistry/hematology analyses showing no indication for adverse reaction. Compared to individual drug treatments, combined therapy resulted in increased efficacy with decreased SARS-CoV-2 shedding and replication early post infection and milder disease as compared to monotherapy.

Dosing for the study was allometrically based on the clinical treatment schedules currently approved for use for COVID-19 treatment. Molnupiravir (MK-4482) is prescribed as an 800mg twice daily oral treatment (1600mg in total) within 5 days of symptom onset [31]. Similarly, nirmatrelvir is an oral treatment prescribed for twice daily oral treatment of 300mg (600mg total) with the addition of 100mg ritonavir (200mg/total) [32]. Each pharmaceutically active compound was assessed in plasma taken at each clinical exam point prior to dosing and in lung homogenates at necropsy to confirm their presence at the desire level and to ensure the absence of any unanticipated drug interaction between the molnupiravir and nirmatrelvir treatments. The levels of each active compound were not negatively affected by the combined treatment regimen.

Phase 3 clinical trials of molnupiravir indicate the drug is effective in preventing severe disease, with lower adverse events documented than the placebo group [10]. Paxlovid™ is also an effective treatment for COVID-19 when administered within 5 days of symptom onset [13]. However, recent reports of viral recrudescence following the cessation of molnupiravir and Paxlovid™ monotherapy in a sub-set of patients has raised the question of efficacy in relation to treatment length and/or dosage [33]. A recent single patient case study indicates that rebound, at least for Paxlovid™, may not correspond to the development of genetic resistance [34]. Combination therapy of MK-4482 and PF-07321332 may counteract the “rebound effect” and thereby enable maintained use of the relatively short 5-day treatment course. Loss of treatment efficacy of a single treatment approach due to viral escape is also always a concern. *In vitro* studies using coronaviruses related to SARS-CoV-2 suggest that molnupiravir may have a high genetic barrier to development of drug resistance [35]. However, recent studies passaging SARS-CoV-2 under suboptimal levels of nirmatrelvir have shown the relatively rapid development of resistance corresponding to defined mutations within the viral protease [36, 37]. The use of a combination therapy as established here would be expected to reduce the possibility for viral escape, as has been established for treatment of other genetically mutable viruses, hepatitis C and human immunodeficiency virus [38, 39].

There are limitations to our study that need to be addressed in future experimental and clinical studies. First, due to the mild-to-moderate level of clinical disease in rhesus macaques following SARS-CoV-2 infection, it was not possible to assess the combination therapy against severe disease. Since SARS-CoV-2 Delta infection likely shows the most severe infection in a NHP model [26], this could be addressed in lethal rodent models or clinical trials. A suggestion of such efficacy against more severe disease is indicated in the recent study using a lethal transgenic COVID-19 mouse model [30]. Second, we only tested the efficacy of the current human dose for monotherapy. Dose reduction might also be achievable in combination therapy which needs to be carefully designed and analyzed. Third, escape mutant development needs to be studied in comparison between mono- and combination therapy, especially given the recent studies suggesting the susceptibility of nirmatrelvir to resistance development *in vitro* [36, 37]. Lastly, clinical trials need to ultimately show the benefit of the combination therapy, which if as tolerable as suggested by our present study, could have benefits such as reducing disease severity in the lower and upper respiratory tract, the latter of which could be a game changer by also reducing transmission.

In conclusion, MK-4482 and PF-07321332 had potent inhibitory effects on SARS-CoV-2 replication, shedding and disease manifestation – now demonstrated in the NHP COVID-19 model, the closest surrogate to humans. Notably, the two drugs were most effective when used in combination. Combination therapy could help to reduce the individual drug doses and slow down drug resistance development. Combination therapy might also improve clinical outcomes in those patients prone towards more severe disease development such as immunocompromised and those with other underlying disease. This study strongly supports the use of MK-4482 or PF-07321332 for the treatment of COVID-19 cases and provides the preclinical evidence for combination therapy as an improved therapeutic option.

## Materials and Methods

### Biosafety and ethics

SARS-CoV-2 studies were approved by the Institutional Biosafety Committee (IBC) and performed in the BSL-4 laboratories at Rocky Mountain Laboratories (RML), NIAID, NIH. IBC-approved standard operating procedures were used for sample removal from biocontainment. RML is an AALAC accredited facility, and the RML Institutional Animal Care and Use Committee approved the animal studies. Animal studies followed institutional guidelines for animal use, the guidelines and basic principles in the NIH Guide for the Care and Use of Laboratory Animals, the Animal Welfare Act, United States Department of Agriculture and the United States Public Health Service Policy on Humane Care and Use of Laboratory Animals. Rhesus macaques were singly housed in adjacent primate cages to enable social interactions between animals. The animal room was climate-controlled with a fixed 12 hours light-dark cycle. Animals were provided commercial monkey chow provided twice daily with vegetables, fruit, and treats to supplement. Water was available *ad libitum*. Human interaction, manipulanda, toys, video and music were provided for enrichment. Animals were monitored for signs of disease at least twice daily throughout the experiment.

### Rhesus macaque study design

Male and female rhesus macaques 2-12 years of age were divided into vehicle (N=5) or treatment (N=5 for MK-4482, N=5 for PF-07321332, and N=5 for the MK-4482/PF-07321332 combination) groups prior to infection. MK-4482 (DC Chemicals), PF-07321332 (DC Chemicals) and ritonavir (DC Chemicals) were first dissolved in DMSO and then resuspended to 5ml total volume in food grade peanut oil for oral delivery. Treated rhesus macaques received either 130mg/kg MK-4482, 20mg/kg PF-07321332 + 6.5mg/kg ritonavir or the combination of all 3 compounds (130mg/kg MK-4482 + 20mg/kg PF-07321332 +6.5mg ritonavir) every 12 hours beginning 12hpi and ending 84hpi. Vehicle control animals received the same treatment volume of peanut oil (5ml) on the same schedule as treatment groups. Animals were infected with a total of 2 × 10^6^ TCID_50_ of the SARS-CoV-2 Delta variant by two routes, intranasal and intratracheal. Intranasal inoculation consisted of a 0.5ml injection directly into each nare (1.0 ml total) with a Mucosal Atomization Device (MAD) system. Intratracheal inoculations were performed with the use of a bronchoscope for deposition of 4mls of virus directly into the main-stem bronchi. All procedures were performed on anesthetized animals. Animals were monitored twice daily and scored blindly every morning by the same person for signs and progression of disease as previously reported [19] In brief, animals were scored based on general and skin/coat appearance, activity, mucosal discharge, respiration, feces and urine output, and appetite. Clinical exams were performed prior to challenge and at D0, D1, D2 and D4. Oropharyngeal, nasal and rectal swabs, as well as whole blood and serum were collected at every exam. Bronchoalveolar lavage samples were collected at D0, D1 and D2, but not at D4 to avoid complications with pathology at time of autopsy. Animals were euthanized on day 4 post-infection and tissues were collected at necropsy for virological and pathology analysis.

### Virus and cells

SARS-CoV-2 variant hCoV-19/USA/KY-CDC-2-4242084/2021 (B.1.617.2, Delta) was kindly contributed by B. Zhou, N. Thornburg and S. Tong (Centers for Disease Control and Prevention, USA). The viral stock was sequenced via Illumina-based deep sequencing to confirm identity and identify any possible contaminants prior to use. Virus propagation was performed in DMEM (Sigma) supplemented with 2% fetal bovine serum (Gibco), 1 mM L-glutamine (Gibco), 50 U/ml penicillin and 50 μg/ml streptomycin (Gibco). Vero E6 cells, kindly provided by R. Baric, University of North Carolina, were maintained in DMEM (Sigma) supplemented with 10% fetal calf serum, 1 mM L-glutamine, 50 U/mL penicillin and 50 μg/mL streptomycin.

### Viral genome detection

A QiaAmp Viral RNA kit was used to extract RNA from swabs or tissue samples (30 mg or less). A Quantifast kit was used according to manufacturer’s protocols and one-step real-time RT-PCR was used to quantify subgenomic viral RNA using a region of the E gene [40]. RNA standards counted by droplet digital PCR were used in 10-fold serial dilutions and run in parallel to calculate viral RNA copies.

### Virus titration assay

To calculate infectious virus titers in tissue samples, tissues were homogenized in 1ml DMEM using a TissueLyzer (Qiagen) and clarified by low-speed centrifugation. Vero-E6 cells were inoculated with 10-fold serial dilutions of homogenized lung or swab samples in 100 µl DMEM (Sigma-Aldrich) supplemented with 2% fetal bovine serum, 1 mM L-glutamine, 50 U/ml penicillin and 50 µg/ml streptomycin. Cells were incubated for six days and then scored for cytopathogenic effects (CPE) and TCID_50_ was calculated via the Reed-Muench formula [41].

### Pharmokinetics

All solvents and extraction reagents were LCMS grade and purchased from Fisher Scientific. Lung sections were extracted and levels of EIDD-1931 were measured as previously described [21]. Plasma samples were processed via dilution of 100 µL of plasma into 300 µl of ice-cold methanol. The precipitant was cleared by centrifugation and the supernatant was prepared for sample injection. Multiple reaction monitoring (MRM) ion pair signals were developed and optimized for ritonavir and PF-07321332 (nirmatrelvir) from standards (Supplemental Table 1). All pharmaceutical actives were measured from a single injection on a Sciex ExionLC™ AC system with a Waters XBridge® Amide column (130□Å, 3.5□µm, 3□mm□×□100□mm) with binary gradient elution from 95% acetonitrile, 0.8% acetic acid, 10□mM ammonium acetate to 50% acetonitrile, 0.8% acetic acid, 10 mM ammonium. All signals were detected in positive mode using a Sciex 5500 QTRAP® mass spectrometer equipped with an electrospray ionization source (CUR: 40, CAD: Med, ISV: 2500, Temp: 450, GS1: 50, GS2: 50). All signals were detected under positive mode ionization and compared to an 8-point standard curve. All molecules of interested were detected using two MRM pairs and quantified using the higher signal to noise MRM. Limit of quantification was 400 pg/ml for PF-07321332 and 300 pg/ml for ritonavir.

### Histopathology

Tissues samples were collected, placed into cassettes and fixed in 10% formalin for 7 days. Prior to removal from BSL-4, tissues were transferred into fresh 10% formalin and incubated an additional 24 hours. Tissues were then processed with a Sakura VIP-6 Tissue Tek, on a 12-hour automated schedule, using a graded series of ethanol, xylene, and PureAffin. Following processing, samples were embedded in Pureaffin paraffin polymer (Cancer Diagnostics, Durham, NC, USA) and sectioned at 5 µm and dried overnight at 42°C prior to hematoxylin and eosin (H&E) staining. IHC tissues were processed using the Discovery Ultra automated stainer (Ventana Medical Systems) with a ChromoMap DAB kit (Roche Tissue Diagnostics cat#760-159) for immunohistochemistry (IHC) staining. Specific immunoreactivity was detected using a validated GenScript U864YFA140-4/CB2093 NP-1 SARS-CoV-2-specific antiserum at a 1:1000 dilution. The secondary antibody was an anti-rabbit IgG polymer (cat# MP-6401) from Vector Laboratories ImPress VR as previously described [21].

### Statistical analyses

Statistical analysis was performed in Prism 9. Difference in viral load and infectious titers between study groups was assessed by ordinary two-way ANOVA (multiple time points) or by multiple comparisons using the Kruskal-Wallace test (single time-points). AUC was calculated and then analyzed by an ordinary one-way ANOVA.

## AUTHOR CONTRIBUTIONS

<u>K. Rosenke, M.A. Jarvis and H. Feldmann</u> contributed to the design, data analysis, and writing of the manuscript.

<u>K. Rosenke, H. Feldmann, J. Lovaglio, M. Lewis, F. Feldmann, W. Bohler, C. Shaia and G. Saturday</u> contributed to the execution of the study

<u>A. Okumura, M.C. Lewis, F. Feldmann, W. Bohler, E. Borhnsen, B. Schwarz, C. Bosio and J. Callison</u> contributed experimental support and data analysis.

<u>G. Saturday and C. Shaia</u> contributed to the pathological analysis.

<u>H. Feldmann and M.A. Jarvis</u> secured funding for the study.

## ACKNOWLEDGMENTS

The authors thank Myndi Holbrook, Craig Martens, Kent Barbian, Stacey Ricklefs and Sarah Anzick of the Genomics Unit of the Research Technology Branch (National Institutes of Allergy and Infectious Diseases (NIAID), National Institutes of Health (NIH)) for their efforts in sequencing viral stocks. The authors also wish to thank the Rocky Mountain Veterinary Branch (RMVB, NIAID, NIH) clinical veterinarians Patrick Hanley and Brian Smith for technical assistance with animal work, Brandon Bailes, Richard Cole, Lydia Crawford, Lisa Heaney, Corey Henderson, Taylor Lippincott, Travis Spencer, Kathy Cordova and Marissa Woods for their efforts in animal care and husbandry, and Tina Thomas, Rebecca Rosenke, and Dan Long for assistance with histology. We also thank Anita Mora, Ryan Kissinger and Austin Athman (Visual and Medical Arts Unit, NIAID, NIH) for help with graphical design. The Delta variant was deposited by the Centers for Disease Control and Prevention and obtained through BEI Resources, NIAID, NIH: hCoV-19/USA/KY-CDC-2-4242084/2021. Vero E6 cells were obtained from Dr. Ralph Baric at the University of North Carolina. The work was mainly funded by the Intramural Research Program of NIAID, NIH and partially through the University of Plymouth.

## DATA AND MATERIALS AVAILABILITY

All data is presented here. Additional information can be requested through the corresponding authors.

### Funding

This study was supported by the Intramural Research Program of NIAID, NIH. This work was part of NIAID’s SARS-CoV-2 Assessment of Viral Evolution (SAVE) Program. Additional funding was provided by the University of Plymouth.

## DISCLAIMER

The opinions, conclusions and recommendations in this report are those of the authors and do not necessarily represent the official positions of the National Institute of Allergy and Infectious Diseases (NIAID) at the National Institutes of Health (NIH), The Vaccine Group Ltd or the University of Plymouth.

---

**Supplemental Table 1:** MRM signals were identified and optimized for ritonavir and PF-07321332 (nirmatrelvir) from standards.Key: MRM: multiple reaction monitoring; DP: declustering potential; EP: entrance potential; CE: collision cell entrance potential; CXP: collision cell exit potential

| Target | MRM pair<br>(m/z) | DP<br>(V) | EP<br>(V) | CE<br>(V) | CXP<br>(V) |
| --- | --- | --- | --- | --- | --- |
| PF-07321332* | 500.0/110.0 | 75 | 10 | 30 | 7 |
| PF-07321332 | 500.0/69.0 | 180 | 5 | 80 | 30 |
| Ritonavir* | 721.0/140.0 | 185 | 15 | 85 | 20 |
| Ritonavir | 721.0/268.0 | 120 | 5 | 30 | 40 |

\*Used for quantification

**Supplementary figure 1.**
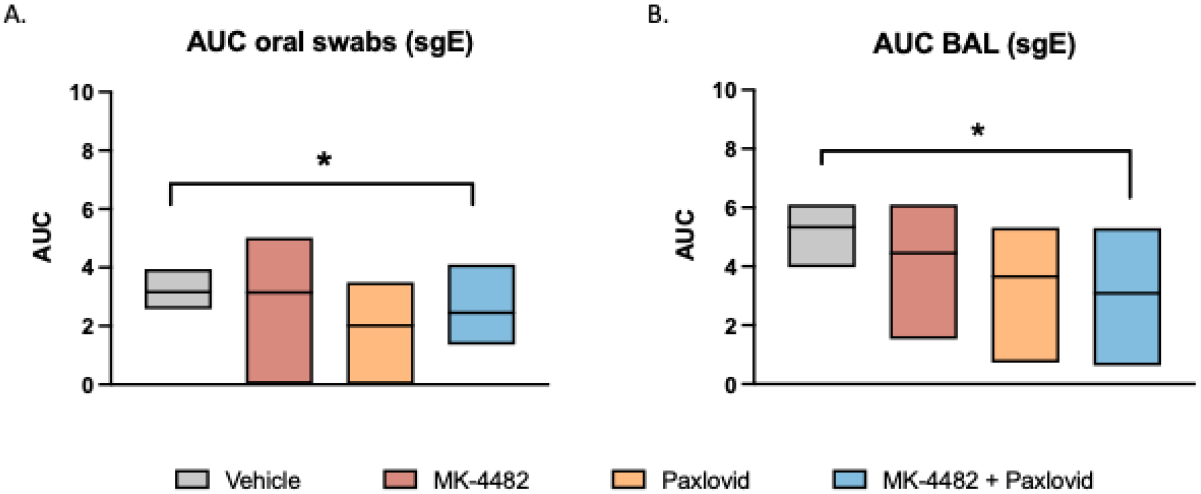
AUC analysis of oral swabs and BAL fluid. Viral RNA loads from oral swabs **(A)** and BAL **(B)** samples were determined by quantitative RT-PCR targeting sgE RNA as a surrogate for replication and shedding. Copy numbers of viral genomes were calculated for each animal per day, AUC was then calculated over the course of the study and displayed in a boxplot with the mean displayed. Ordinary one-way ANOVA with multiple comparisons were used to evaluate significance (*P-value = 0.01 to 0.05).

**Supplementary figure 2.**
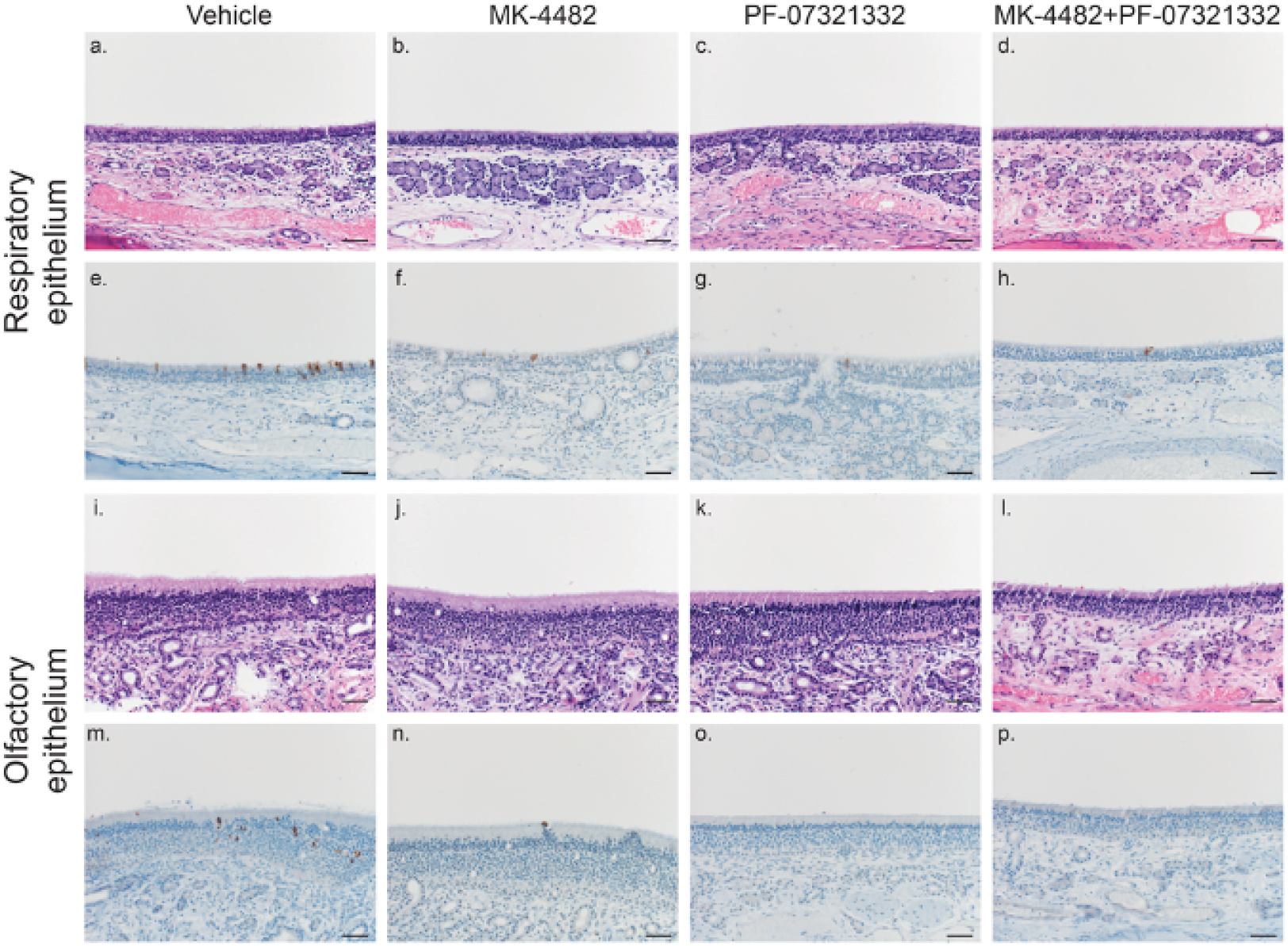
Combination therapy reduced antigen load in olfactory and respiratory epithelium from nasal turbinates in SARS-CoV-2 infected rhesus macaques. Tissues were collected on 4dpi and stained with H&E or IHC for analysis. **H&E staining of representative tissues sections of the respiratory epithelium (A-D)**. No pathology was found in H&E stains of the nasal turbinates. **IHC staining of SARS-CoV-2 antigen in respiratory epithelium (E-H)**. Reduced or no IHC stain was found in the respiratory epithelium of MK-4482, PF-07321332 and MK-4482 + PF-07321332 treated animals compared to vehicle controls. **H&E staining of representative tissues sections of the olfactory epithelium from nasal turbinate (I-L)**. No pathology was found in the olfactory epithelium. **IHC staining of SARS-CoV-2 antigen in olfactory epithelium (M-P)**. IHC analysis did show scattered immunoreactivity in the vehicle treated animals and little to no observable viral antigen in the MK-4482, PF-07321332 and MK-4482 + PF-07321332 treated animals 200X, Bar=50µm

